# Modeling Pulse Dynamics of Juvenile Fish Enables the Short-term Forecasting of Population Dynamics in Japanese Pufferfish: A Latent Variable Approach

**DOI:** 10.1101/2022.01.26.477932

**Authors:** Shota Nishijima, Shigenori Suzuki, Ryo Fukuta, Makoto Okada

## Abstract

The time lag between data collection and management implementation is a source of uncertainty and bias in the calculation of acceptable biological catch. Here, we developed a method to use small-scale survey data for juvenile Japanese pufferfish (*Takifugu rubripes*) to shorten this time lag and achieve accurate short-term forecasting. A survey of juvenile fish at a local sandy beach in Ise-Mikawa Bay, Japan provides data for the strength of year classes before fisheries recruitment; however, it is difficult to use the raw data owing to the small sample size and large observation errors. We found that a random-effect model overcame these problems and more accurately predicted pulse patterns of catch rates to derive a standardized recruitment index compared with a fixed-effect model. We then showed that a stock assessment model using the standardized recruitment index from the random-effect model outperformed models without the standardized recruitment index with respect to retrospective forecasting ability. This study highlights the applicability of a latent-variable approach for standardizing small-scale survey data and thereby for unbiased forecasting of short-term fish dynamics.

## Introduction

Future projections of fish population dynamics are generally difficult but are necessary for stock assessment and the management of fisheries resources (Beveren et al. 2021). An objective of stock assessment is to recommend the acceptable biological catch (ABC) in the near future under a harvest strategy considering the long-term sustainable use of a certain fisheries stock (Wiedenmann et al. 2016). Because stock assessment models usually estimate abundance and fishing mortality rates only for the period when data are available, fisheries scientists need to forecast population dynamics of the fisheries stock for at least a few years up to the ABC year (Cadima 2003). However, estimates of the stock status in the most recent years show high uncertainty and bias, and future forecasting can increase this uncertainty (Deroba and Bence 2008; Punt et al. 2016). In addition, the prediction of fisheries recruitment is challenging owing to the high variability (Beveren et al. 2021). The inevitable time lag between data collection and management implementation requires the short-term forecasting of fish dynamics and can be a source of uncertainty and bias in ABC, thereby driving overfishing and the under-use of fisheries resources (MacKenzie et al. 2008; Li et al. 2016).

Extensive efforts have focused on the development of methods for accurate future forecasting and the evaluation of the importance of filling in the data management time lag. A potential approach to improve future forecasting is to use a climatic or environmental variable for which recent values are available (Costello et al. 1998; MacKenzie et al. 2008). Previous studies have shown, however, that incorporating environmental factors into stock assessment and short-term forecasting is unlikely to be successful because long, complex life histories before fisheries recruitment (Haltuch et al. 2019) make the relationship between recruitment and environments ambiguous, which does not substantially improve the ability to reach management goals (Punt et al. 2014). Although surveys to determine the strength of a year class before fishery recruitment can provide direct prior information on the strength of the year class in the current year (Hashimoto et al. 2019), the feasibility of this approach depends on the costs and human resources required for the surveys. A few studies have shown that a reduced frequency of surveys and thereby an increased time lag between data collection and management efforts did not substantially affect the performance of stock assessments, and the costs of surveys exceeded the societal benefits generated from fisheries for some stocks (Zimmermann and Enberg 2017; Hutniczak et al. 2019). Accordingly, it is necessary to develop an approach that improves the performance of stock assessments and future forecasting based on low-cost surveys, especially for minor fish stocks for which exhaustive surveys are difficult to conduct.

In the Ise-Mikawa Bay stock of Japanese pufferfish (*Takifugu rubripes*) ABC has been overestimated in recent years (Suzuki et al. 2021). In Japan, annual stock assessments on domestic fisheries resources usually use data up to the previous fishing year and calculate ABC for the next fishing year (Okamura et al. 2020). There is, thus, a two-year time lag between data collection and management implementation. A longline fishery is the largest fishery of this pufferfish stock and harvests mainly age 1 fish (Nishijima et al. 2019; Suzuki et al. 2021). However, catch data for this cohort in this ABC year are not available at the time of stock assessment due to the two-year lag. The number of recruits in the future is assumed to be the five-year average because the stock-recruitment relationship is ambiguous; however, this prediction tends to be positively biased because the number of recruits has decreased gradually in recent years (Suzuki et al. 2021). In other words, the combination of fishery selectivity towards young ages and the biased prediction of recruitment has caused the overestimation of stock biomass and ABC in the near future. Because the stock has long been kept at low levels and is overfished (Suzuki et al. 2021), the overestimation of future stock abundance and ABC is a serious issue that jeopardizes the sustainability of pufferfish fisheries.

A small-scale survey has been conducted every year since 2004 to monitor the recruitment of this pufferfish stock before they are caught by fisheries. This survey samples juvenile fish that are transported to the Shiroko seacoast of Suzuka in Mie Prefecture from May to July (Fig. 1). The survey data can provide timely information on the number of age 0 fish in the current year and are available for the meeting of the annual stock assessment in August. However, these raw data are difficult to use directly owing to the small sample size, large observation errors, and irregular sampling frequency. The relationship between survey catch-per-unit-effort (CPUE) and date exhibits a pulse pattern with the highest peak at an intermediate point because juvenile fish gradually arrive at this coastal sandy area and then leave after feeding and growing; however, this pulse pattern was unclear for some years (Fig. 2). Furthermore, there were no captures over five sampling days in 2017, meaning that the nominal CPUE was zero in that year (Fig. 2). This highlights the large observation errors and the need for standardization.

**Figure 1:**
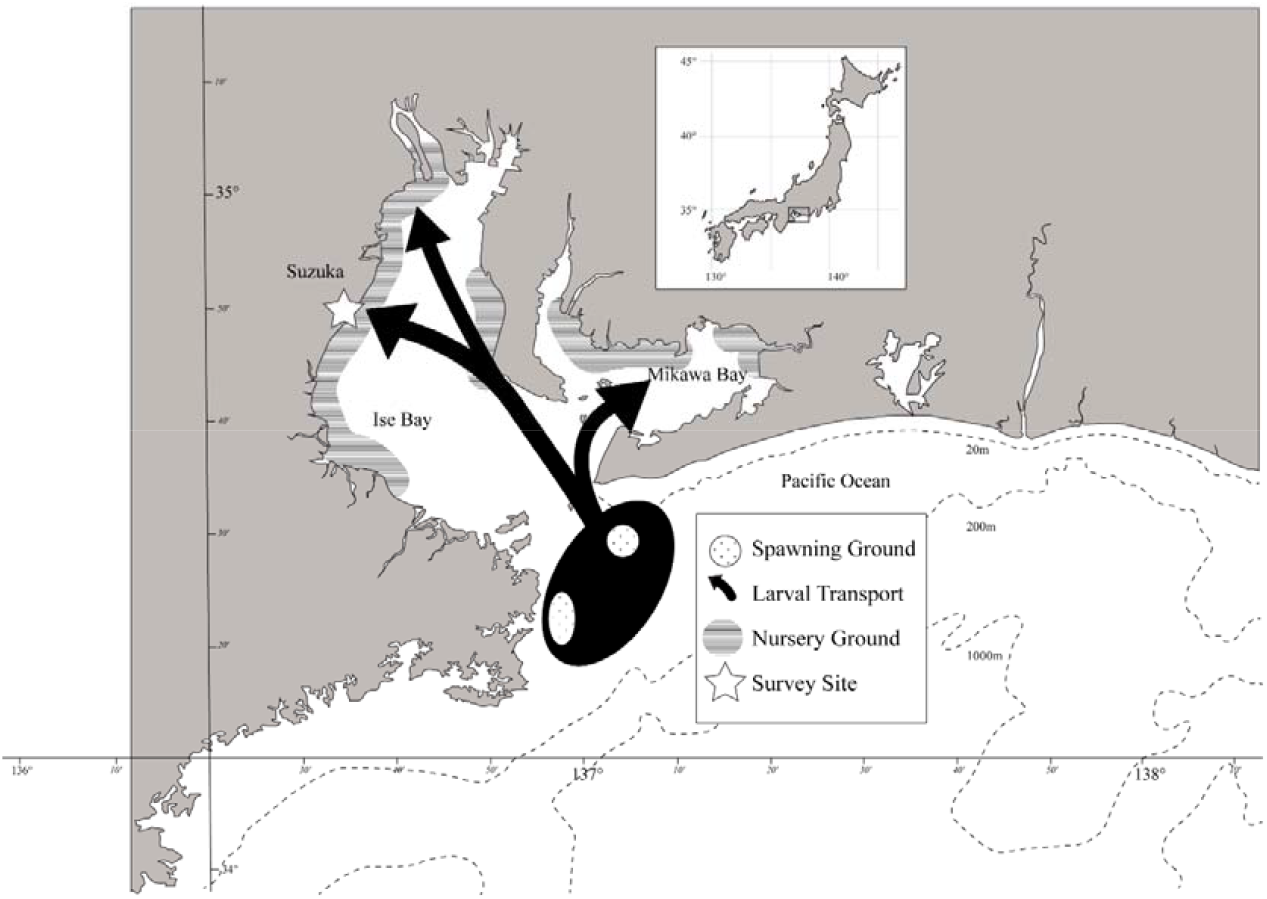
Life history from hatching to juvenile stages of Japanese pufferfish in Ise-Mikawa Bay and the location of the surf-net survey at Suzuka, Mie prefecture, Japan.

**Figure 2:**
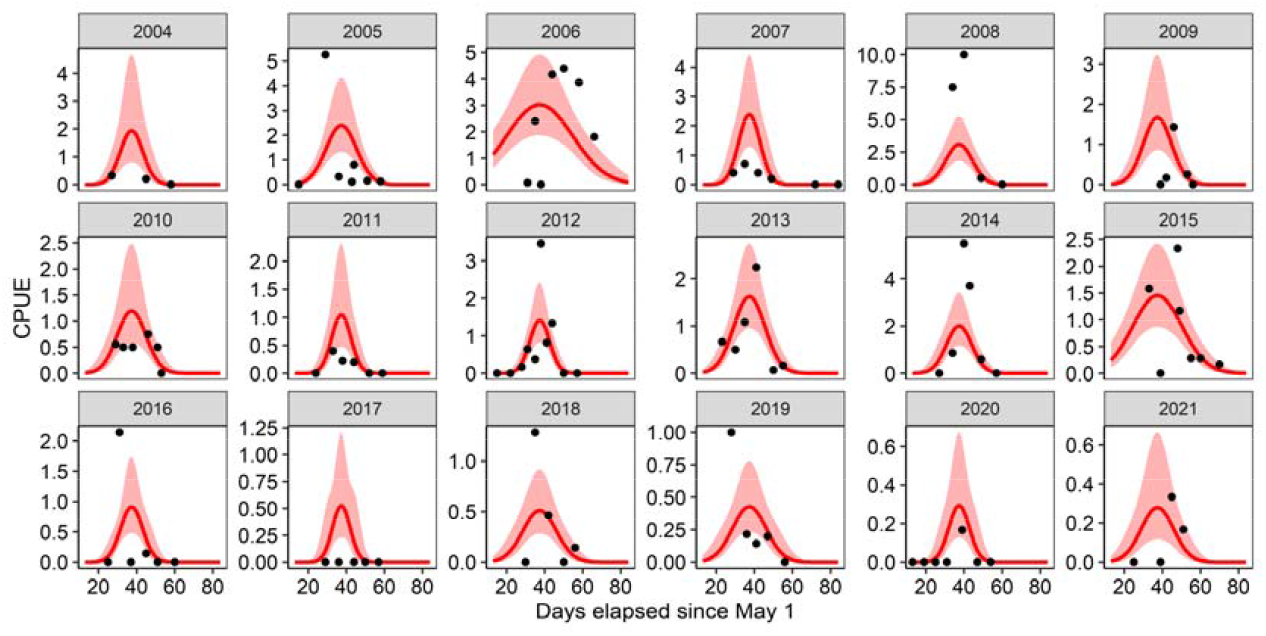
Relationship between CPUE (individuals/tow) and days elapsed since May 1 from 2004 to 2021 (black dots: observation, red lines: prediction by the best model, red shadows: 80% confidence intervals).

In this paper, we develop a latent variable approach for CPUE standardization to accurately forecast short-term population dynamics of Japanese pufferfish based on survey data. Random effects models with latent variables are prevailing in recent fisheries science because they can robustly estimate a large number of parameters, which would cause overfitting or a failure to converge if using fixed effects (Nielsen and Berg 2014; Thorson and Barnett 2017; Nishijima et al. 2021). Here, we aim to stably estimate appropriate positive values of expected CPUE, even for zero-catch samples, by using a random effects model. Our model describes pulse patterns of CPUE as a quadratic function and estimates interannual variation in pulse dynamics as fixed effects or random effects. We first compare the predictive abilities of fixed-effect and random-effect models and derive a standardized recruitment index from the model with highest predictive ability. We then conduct ‘retrospective forecasting’ or ‘hindcasting’ (Brooks and Legault 2016; Kell et al. 2016) using a virtual population analysis (VPA) for the stock assessment of Japanese pufferfish (Suzuki et al. 2021) and compare the performance of short-term forecasting among VPA models with and without the standardized recruitment index. Retrospective forecasting using the VPA method provides short-term forecasts using truncated assessment data (data for the latest year(s) are removed sequentially); accordingly, we compare results obtained with truncated data and with full time series data (Brooks and Legault 2016). We illustrate that even small-scale survey data can be standardized by a latent-variable model, shortening the data management time lag and improving the short-term forecasting of pufferfish population dynamics.

## Materials and Methods

### Life history and surf-net survey

The Japanese pufferfish spawns in limited areas just outside of the entrance of Ise-Mikawa Bay for a short period of time from April to May (Fig. 1). After hatching, larvae are passively transported into Ise-Mikawa Bay and reach sandy coasts with a total length of approximately 10 mm (Nakajima et al. 2008b; Tsumoto 2013; Okada et al. 2015; Suzuki et al. 2015). Juvenile fish then remain in the coastal sandy zones as nursery areas from late June to July, growing to a total length of approximately 30 mm, and gradually migrate to shallow areas with a depth of ≤10 m inside Ise-Mikawa Bay (Suzuki et al. 2021). Young-of-the year fish (YOY) are caught by small-scale trawl fisheries inside and outside of the bay beginning in the autumn.

A local surf-net survey has been conducted at the Shiroko seacoast of Suzuka, Mie Prefecture (Fig. 2) every year since 2004 to monitor the recruitment of Japanese pufferfish. This site was selected because many juvenile fish (10 mm) could be efficiently sampled by surf-nets (Nakajima et al. 2008b). The surf-net survey is conducted about once a week from late May to early July (5.83 ± 1.54 [SD] days/year). The sample size (i.e., the total number of days over 18 years) was *N* = 105. The number of tows per day (effort size) was 7.30 ± 3.03, on average, and the distance of tows was fixed at 50 m. This small-scale survey is usually performed by two people on behalf of Mie Prefecture Fisheries Research Institute and, therefore, is a relatively low-cost method. In a few years at the beginning of the survey, we used different kinds of surf-nets from the current surf-net (Table A1; Nakajima et al. 2008b, Tsumoto 2013, Suzuki et al. 2015). Because the size of surf-nets may affect catchability, we compared three cases using the best model (see below): (1) adjusting effort by the area of wind nets, (2) adjusting effort by the length of wind nets, and (3) no adjustment. We found that case (2) had a slightly better fit (i.e., higher likelihood) than the other two cases. Therefore, we show results of case (2), which was adjusted effort by the length of wind nets. We defined a single tow with the small seine net since 2008 as the per-unit effort.

The annual stock assessment in Ise-Mikawa Bay is conducted in August. Thus, the surf-net survey of the juvenile stage of YOY fish from May to July provides important prior information, before fisheries recruitment, which can be used as the most recent data in the annual stock assessment.

### Model development

CPUE showed a pulse pattern with a peak at an intermediate date (Fig. 2). The limited spawning locations and spawning period, passive movement of larvae, and transient utilization of nursery grounds, explained in the above subsection, result in high juvenile catch rates in the surf-net survey for only a short time period. We described this pulse pattern using a quadratic function of days elapsed since May 1st (*t*) in a year (*y*):

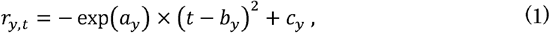

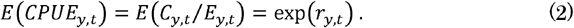

Here, *r_y,t_* is the log-transformed expected CPUE value on day *t* in year *y*. *a_y_* controls the shape or width of pulse in year *y* (i.e., a larger *a_y_* indicates a sharper shape or narrower pulse width). We used the exponential function of *a_y_* restricted to concave-down. *b_y_* and *c_y_* represent the timing and size, respectively, of the pulse peak. The function of expected CPUE is identical to the Gaussian function used to represent a variety of natural phenomena. Therefore, our mathematical formulation is a simple approach to express the pulse dynamics. The observed catch number (*C_y,t_*) is an integer that is greater than or equal to zero. We used a negative binomial distribution to account for overdispersion resulting from large observation errors. Although the data included many zero values (39 of 105 samples), a model diagnostic showed that a zero-inflated model is not necessary (Fig. S1f in Online Supplementary Material). We defined the expected catch number *μ_y,t_* = *E*(*C_y,t_*) as the product of effort (number of tows adjusted by the length of wind nets, see above and table A1) and expected CPUE (i.e., *μ_yt_* = *E_y,t_* × exp(*r_y,t_*)). The variance in catch number is *μ_y,t_* × (1 + *φ* × *μ_y,t_*, where the *φ* parameter represents the degree of overdispersion (*φ* > 0).

We consider five types of interannual variation in *a_y_*, *b_y_*, and *c_y_*. These five types are used to estimate year effects by the vector autoregressive spatio-temporal (VAST) model for CPUE standardization (Thorson 2019). Below, we show these types using *c_y_* as an example; the same types apply to *a_y_* and *b_y_*.

1. Fixed: *c_y_* is estimated by fixed effects,
2. Constant: no interannual variation (*c_y_* = *c*),
3. White noise: *c_y_* is estimated independently in each year by random effects (*c_y_*~Normal(c, 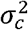)),
4. AR(1): autocorrelation between a year and the previous year (cor(*c_y_*,*c_y–1_*) = *ρ_c_*), and
5. Random walk: a random value is obtained depending on the value in the previous year (*c_y_*~Normal(*c_y–1_*, 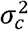)).

We estimated the parameter in each year (*c_y_*) as fixed effects for types 1 and 2 and as random effects by a latent variable model for types 3, 4, and 5. The parameters *c*, *ρ_c_*, and *σ_c_* were estimated as fixed effects. Type 3 assumes temporal independence between any two years, while types 4 and 5 assume temporal autocorrelation between two successive years. We considered autocorrelation here because influential oceanographic effects might result in autocorrelation. The latent variables are inferred under the restriction of estimated variance; therefore, using random effects generally resulted in less fluctuation than that using fixed effects. Importantly, this feature makes it possible to estimate a valid value for a year when the total catch throughout the survey season is zero (e.g., 2017; Fig. 2). In other words, when using fixed effects (type 1 above), *c_y_* is estimated as a highly negative value and the expected CPUE becomes essentially zero for such a year, whereas this problem is avoided by using random effects. We estimated the fixed and random parameters by using the R package ‘TMB’ (Kristensen et al. 2016). TMB is an abbreviation for “template model builder,” which enables the fast computation of complex random effect models with an automatic differentiation algorithm and the Laplace approximation via the maximum (marginal) likelihood method.

We considered the five types of variation for the three parameters and thus analyzed 125 (=5^3^) models in total. We selected the best model by the following steps. First, we excluded models that did not converge or showed Hessian matrix errors (62 models). Second, we selected the two models with the lowest Akaike’s information criterion (AIC) or Bayesian information criterion (BIC) values. Because the penalty per estimated parameter is larger for BIC than for AIC, BIC favors a more parsimonious model. Lastly, we performed leave-one-out cross validation (LOOCV) to compare model performance. We estimated parameters from the data when excluding one sample and calculated the predictive error from the excluded sample. We used negative log-likelihood values as the measure of predictive error, following Thorson and Barnett (2017). We replicated the calculation of predictive error for each sample (*N* = 105) and selected the model with the lower mean predictive error as the best model. We conducted a simulation-based model diagnostic (Hartig 2020) to check model assumptions and residuals for the best model. Methods and model diagnostic results are shown in the Online Supplementary Material.

### Derivation of the abundance index

The predicted CPUE can be analytically integrated using the integral of the Gaussian function:

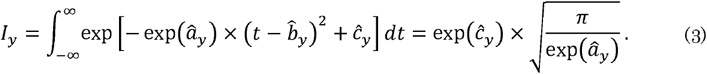

This value is the area surrounded by the predicted curve and *y* = 0 (Fig. 2) and corresponds to the expected total catch when the sampling survey is conducted with per-unit effort (one tow) every day. Integrating the elapsed days *t* between negative and positive infinities may seem odd because the elapsed days should be finite. However, this assumption has a negligible impact because predicted CPUEs are close to zero outside of the survey period (Fig. 2). We therefore used this value as the index of year *y*. It is noteworthy that equation (3) shows that an index value is most sensitive to changes in peak size (*c_y_*), followed by pulse width (*a_y_*), while the peak timing (*b_y_*) has no impact on the index value. We applied a generic method for bias correction in the random effects model to the computation of index values (Thorson and Kristensen 2016). We calculated confidence intervals from the delta method implemented in TMB.

### Application to the tuned VPA and retrospective forecasting

We used an abundance index for naturally occurring YOY (hereafter referred to as the age-0 index for simplicity), developed here, in a tuned VPA to investigate its effect on stock assessment and short-term forecasting. We analyzed four models: (1) without the age-0 index (w/o age-0 index), (2) with the age-0 index using nominal max CPUE (nominal max), (3) with the age-0 index using nominal mean CPUE (nominal mean), and (4) with the standardized age-0 index obtained from the above procedure (standardized). The model without the age-0 index corresponds to the current stock assessment model used in Japan (Suzuki et al. 2021). The nominal max and mean indices cannot be used directly because the total catch was zero in 2017 and thereby cannot be log-transformed. We added the minimum positive value (i.e., the minimum except for 2017) as a small constant to the values for all years. Although we analyzed the case when the small constant is the half of the minimum positive value, we do not show this case because the fitting of this index was worse than that for the case when the small constant is the minimum positive value.

We used the same data that were used for the annual stock assessment (Suzuki et al. 2022): catch-at-age, weight-at-age, and maturity-at-age are available from ages 0 to 3+ years from the fishing years 1993 to 2020 and the natural mortality coefficient was set to M = 0.25, estimated from the generation time of this species. In addition to an age-0 index, an abundance index for this stock for age 1 has been used from fishing year 1995 to 2020 and was obtained by applying the DeLury method to the CPUE of longline fisheries, accounting for time-varying catchability (Nishijima et al. 2019). We deterministically calculated numbers-at-age backward from the catch-at-age and natural mortality in the tuned VPA. We used Pope’s approximation (Pope 1972) in this backward process and age-specific fishing mortality coefficients (*F*). We assumed that *F* at age 2 in each year was equal to that at age 3+ in each year.

We estimated *F* in the terminal year (2020) by separating the age classes into two groups, 0 and 1–3+, because fish at age 0 were caught by small-scale trawl fisheries, whereas fish older than age 0 were caught mainly by longline fisheries (Suzuki et al. 2021). We assumed that the selectivity between ages 1 and 3+ in the terminal year was equal to that averaged over the last three years and used the age-1 index to estimate the terminal *F*. For the model without the age-0 index (w/o the age-0 index), we assumed that *F* at age 0 was the average value over the last two years by following the annual stock assessment (Suzuki et al. 2021): 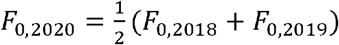. The negative log-likelihood to be minimized in this case is as follows:

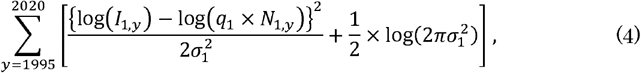

where *I*_1,*y*_ and *N*_1,*y*_ are the age-1 index and the number of age-1 fish, respectively, in year *y*, *q*_1_ is a proportionality constant, and σ_1_ represents the measurement error for the age-1 index. However, for the models with an age-0 index (nominal max, nominal mean, and standardized), we estimated *F* at age 0 in the terminal year by relaxing the assumption in the model without the age-0 index. It is to be noted that these age-0 indices derived from the surf-net survey do not target all recruited fish but only wild YOY fish; the pufferfish stock included the wild population as well as individuals released from hatchery production, and the indices represent only the former because the surf-net survey samples only wild juvenile fish (Nakajima et al. 2008a). Therefore, the number of recruits from hatchery-reared fish must be excluded from the total number of recruits in fitting an age-0 index in the tuned VPA. The number of recruits from hatchery-reared fish was calculated by an additive probability estimated from a mark-recapture experiment (Suzuki et al. 2022). We then calculated the number of recruits from wild fish by subtracting the estimated number of recruits from hatchery-reared fish 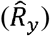 from the total number of recruits (*N*_0,*y*_) when fitting the age-1 indices to the tuned VPA. The negative log-likelihood is expressed as follows:

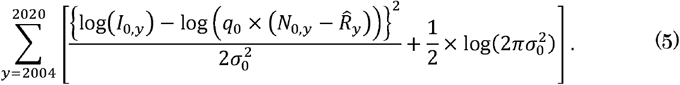

where *I*_0,*y*_ is the value of the age-0 index in year *y*, *q*_0_ is a proportionality constant, and *σ_0_* represents the magnitude of the measurement error for the age-0 index. We estimated parameters so that the sum of negative log-likelihoods in equations 4 and 5 were minimized.

These age-0 indices are available one year ahead of the terminal year in the tuned VPA (i.e., the value in 2021 is available, although the tuned VPA can use only values up to 2020), enabling the short-term forecasting of the number of recruits from wild fish. In the model without the age-0 index (as in the annual stock assessment; Suzuki et al. 2021), we used the average number of recruits in the last 5 years as the predicted recruitment in 2021: 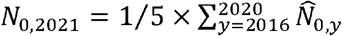. By contrast, we could predict the number of recruits from wild fish using the age-1 index for 2021: *N*_0,2021_ – *R*_2021_ = *I*_0,2021_/*q*_0_. Because the number of recruits from hatchery-reared fish in 2021 (*R*_2021_) was impossible to obtain at present, we averaged the estimated number of recruits from hatchery-reared fish in the last 5 years: 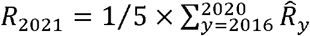. We used the sum of the predicted numbers of recruits from wild fish and hatchery-reared fish as the number of recruits in 2021.

We calculated the numbers of fish of other ages in 2021 by forward calculation from the number and *F* at age in the terminal year 2020. We also predicted the catch in 2021 by assuming that *F* at age in 2021 was equal to the average of the last 3 years (2018–2020): 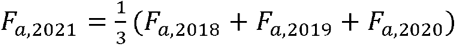. This process of short-term future prediction follows the method used for the Japanese stock assessment.

To evaluate the short-term future prediction performance, we performed a retrospective analysis with future projections, called ‘retrospective forecasting’ or ‘hindcasting,’ in which we sequentially removed the data for the latest year(s) and estimated and forecasted the stock status up to one year ahead in the same way (Brooks and Legault 2016; Kell et al. 2016; Okamura et al. 2021). We compared the forecasted stock status based on the truncated data with the estimated value based on the full data using Mohn’s rho (Mohn 1999):

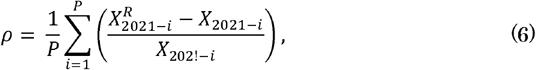

where *X*_2021-*i*_ is the estimate for year *X*_2021-*i*_ using the full data and 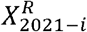 is the corresponding value using the truncated data. We used (1) the number of recruits, (2) total stock biomass, (3) spawning stock biomass (SSB), (4) average *F*, and (5) catch biomass as measures of retrospective forecasting bias. We removed 10 years at maximum (*P* = 10).

## Results

### Model selection

Models selected by AIC and BIC were different. The top models selected by AIC estimated the annual variation in peak size (*c_y_*) as fixed effects (Table 1). The model with minimum AIC estimated the annual variation in peak size (*c_y_*) as fixed effects and peak timing (*b_y_*) as random effects (white noise) and assumed a constant pulse width (*a_y_*).

**Table 1:** Top five models selected by either AIC or BIC.

| Rank | df | Log-likelihood | ΔAIC or ΔBIC | Pattern of annual variation* | | | Cross-validation error rate (mean ± SD) |
| --- | --- | --- | --- | --- | --- | --- | --- |
| | | | | $a_y$ | $b_y$ | $c_y$ | |
| AIC |  |  |  |  |  |  |  |
| 1 | 22 | -231.95 | 0 | Const | WN | FE | 3.14 ± 4.75 |
| 2 | 24 | -230.02 | 0.13 | AR(1) | WN | FE |  |
| 3 | 22 | -232.3 | 0.69 | WN | Const | FE |  |
| 4 | 23 | -231.5 | 1.1 | Const | AR(1) | FE |  |
| 5 | 23 | -231.61 | 1.32 | AR(1) | Const | FE |  |
| BIC |  |  |  |  |  |  |  |
| 1 | 5 | -253.26 | 0 | WN | Const | RW | 2.44 ± 2.03 |
| 2 | 5 | -254.78 | 3.04 | WN | RW | RW |  |
| 3 | 4 | -257.29 | 3.41 | RW | Const | RW |  |
| 4 | 6 | -253.15 | 4.43 | AR(1) | Const | RW |  |
| 5 | 6 | -253.26 | 4.65 | WN | WN | RW |  |

Note: FE, fixed effect; Const, constant; WN, white noise; AR(1), first-order autoregressive; RW, random walk. Detailed definitions are provided in the *Model* *development* subsection in the Materials and Methods.

Models selected by BIC estimated the annual variation in pulse width (*a_y_*) and peak size (*c_y_*) as random effects (Table 1). The top models except for models two and five assumed a constant pulse timing peak timing (*b_y_*). The model with the minimum BIC estimated the annual variation in peak size (*c_y_*) by a random walk and pulse width (*a_y_*) as random effects (white noise) and assumed a constant peak timing (*b_y_*). Moreover, the model with the minimum BIC had also the minimum AICc.

In LOOCV, the model with the minimum BIC had a lower predictive error (negative log-likelihood) than that of the model with the minimum AIC (Table 1). This suggests that a random effects model of annual variation in peak size (*c_y_*) has a higher predictive ability than that of a fixed effect model. Based on this result, we used the model with the minimum BIC as the best model in the following analyses.

### Temporal trends of parameters and index

The best model revealed a random fluctuation in pulse width (Fig. 3a), showing that the duration of pulsed CPUE was longer in 2006 and 2015 than in other years (Fig. 2). The peak timing was constant among years and was detected at around June 7 (Fig. 3b). The peak size, estimated by a random walk, exhibited a decreasing trend since 2014 (Fig. 3c).

**Figure 3:**
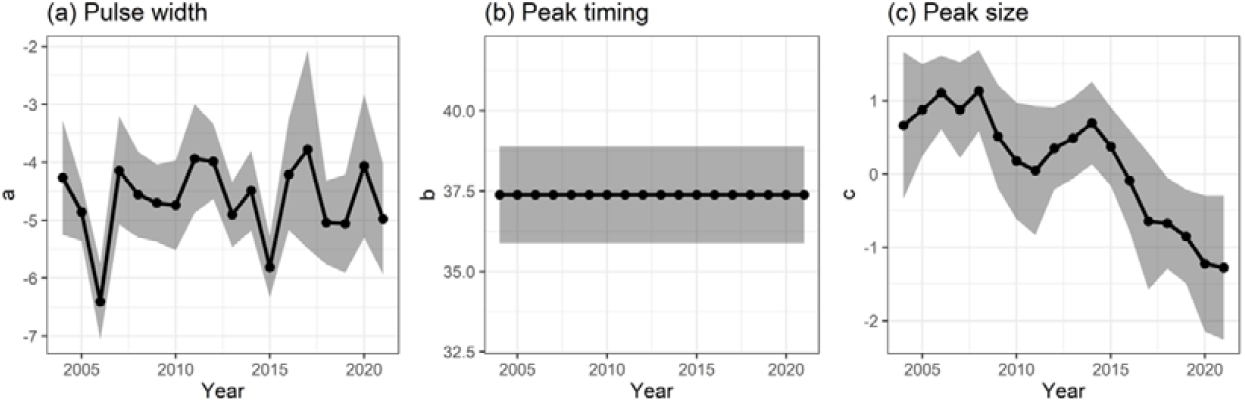
Yearly estimates of pulse width (a), peak timing (b), and peak size (c) with 80% confidence intervals in the best model.

The temporal trend in the standardized abundance index was smoother than those of the nominal mean index and nominal max index (Fig. 4). In 2006, a higher value was obtained for the standardized index than for the nominal indices, whereas the opposite was true in 2008 and 2014. Importantly, a positive value was estimated for 2017, when the total catch was zero (Figs. 2, 4). This is because random-effect models can estimate a finite value for log-transformed expected catch numbers from the estimated mean and variance and attribute zero values to observation errors.

**Figure 4:**
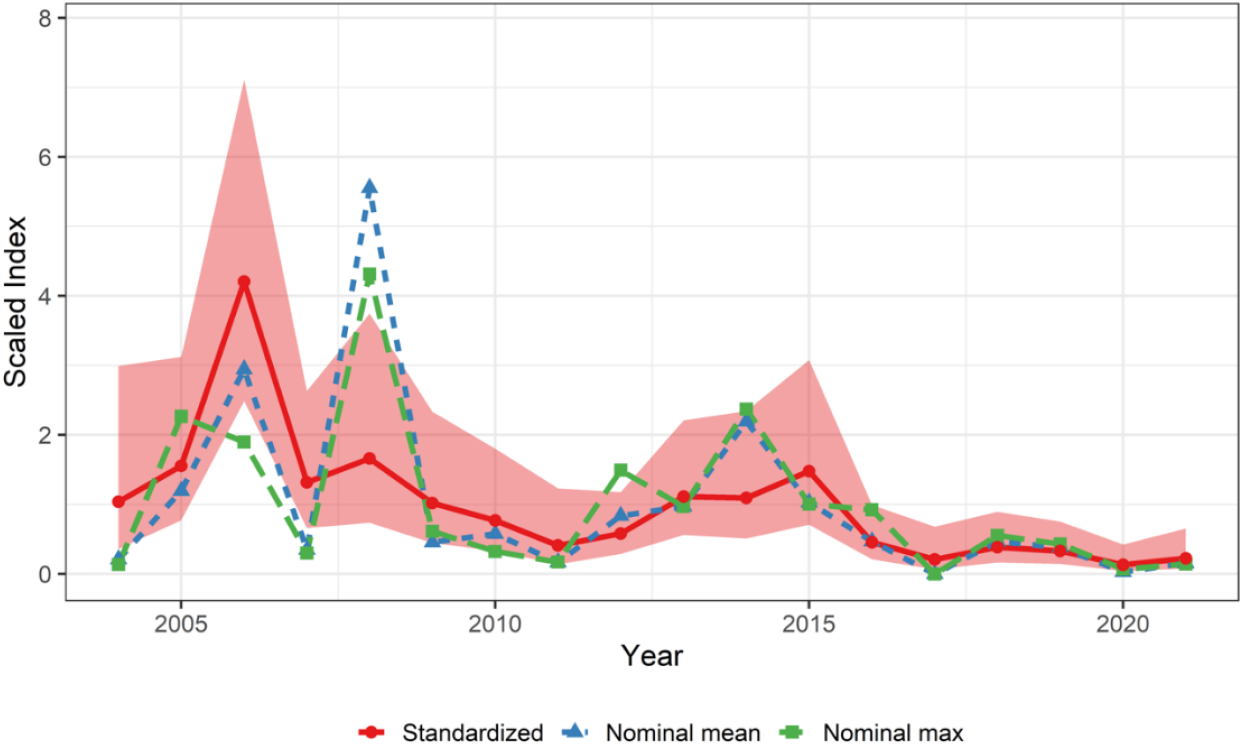
Temporal trends in the standardized index (red solid line), nominal mean index (blue dotted line), and nominal max index (green dashed line). Red shadows indicate 80% confidence intervals. The index values were scaled by their averages for comparison.

### Application to the tuned VPA and retrospective forecasting

The forecasted number of recruits in 2021 was higher in the VPA without the age-0 index than in the tuned VPAs with age-0 indices; however, there were no substantial differences among the models with the standardized and nominal indices (Fig. 5). The estimates of total biomass and spawning stock biomass from 2019 to 2021 were lower in the model with the standardized index than in the other models. This is because the model with the standardized index estimated the lowest number of recruits in 2018 and 2019 and the highest fishing mortality rate since 2018. The predicted catch biomass in 2021 did not differ substantially among models. The standard deviation in the fitting of the abundance index was much smaller for the standardized index (σ_0_= 0.45) than those for the nominal indices (nominal max: σ_0_= 0.87, nominal mean: σ_0_= 0.70), showing a better fitting of the standardized index (Fig. S3 in Online Supplementary Material).

**Figure 5:**
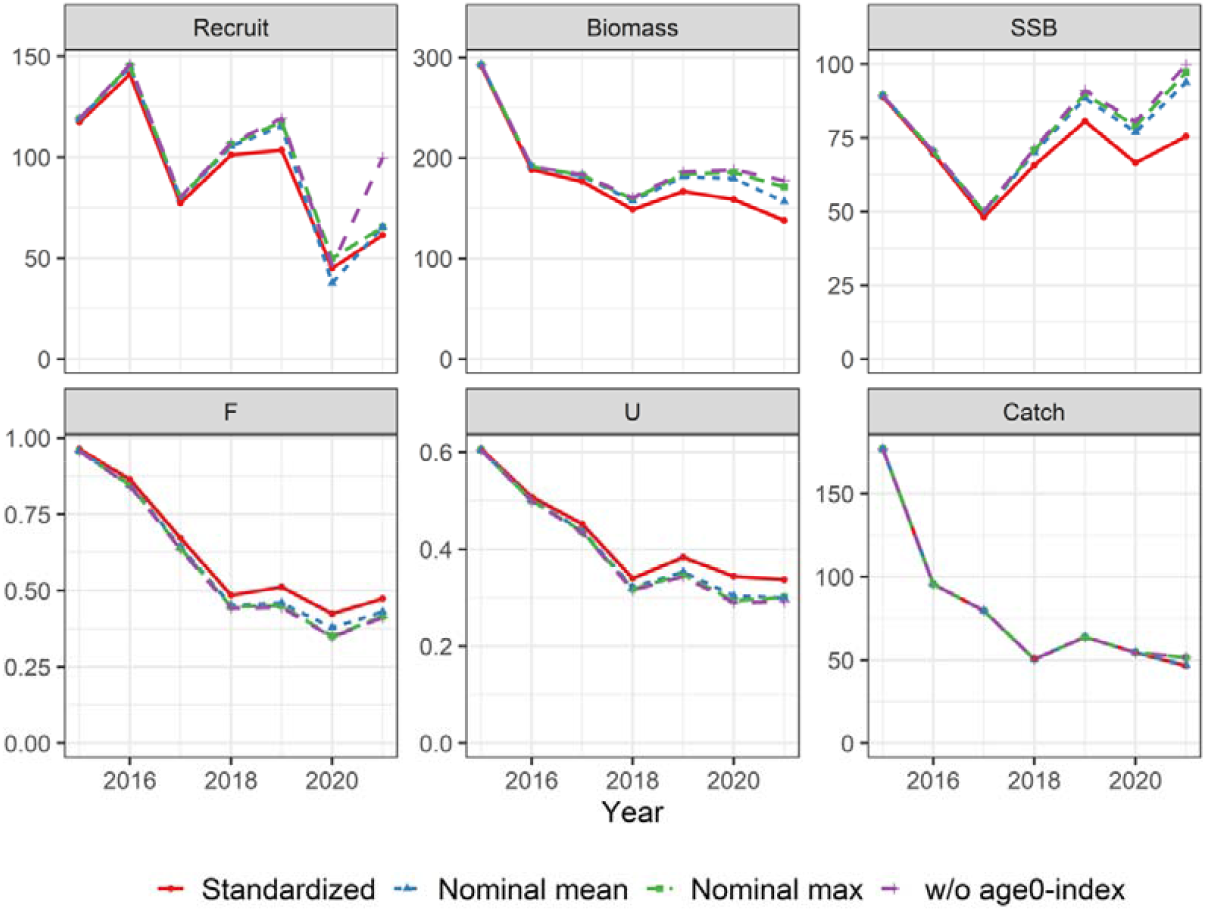
Recent estimates of the number of recruits (left upper), total biomass (middle upper), spawning stock biomass (right upper), average *F* (left lower), exploitation rate (catch biomass divided by total biomass) (middle lower), and catch biomass (right lower) in VPA with the standardized index (red solid lines), nominal mean index (blue dotted lines), nominal max index (green dashed lines), and with no age-0 index (purple long-dashed lines). The values in 2021 are obtained by one-year-ahead forecasting.

The retrospective forecasting analysis showed overestimation in recruitment, total biomass, SSB, and catch biomass in the models with no age-0 index and with the nominal indices (Fig. 6). The biases were highly reduced in the model with the standardized index. Overestimation bias in SSB was still found in the model with the standardized index; however, it was clearly improved compared with those of other models. Higher values for total biomass and SSB in 2021 were forecasted by VPA with no age-0 and nominal indices than by VPA with the standardized index (Fig. 5) and would be more likely to be overestimated. All models showed low bias in fishing mortality coefficients.

**Figure 6.**
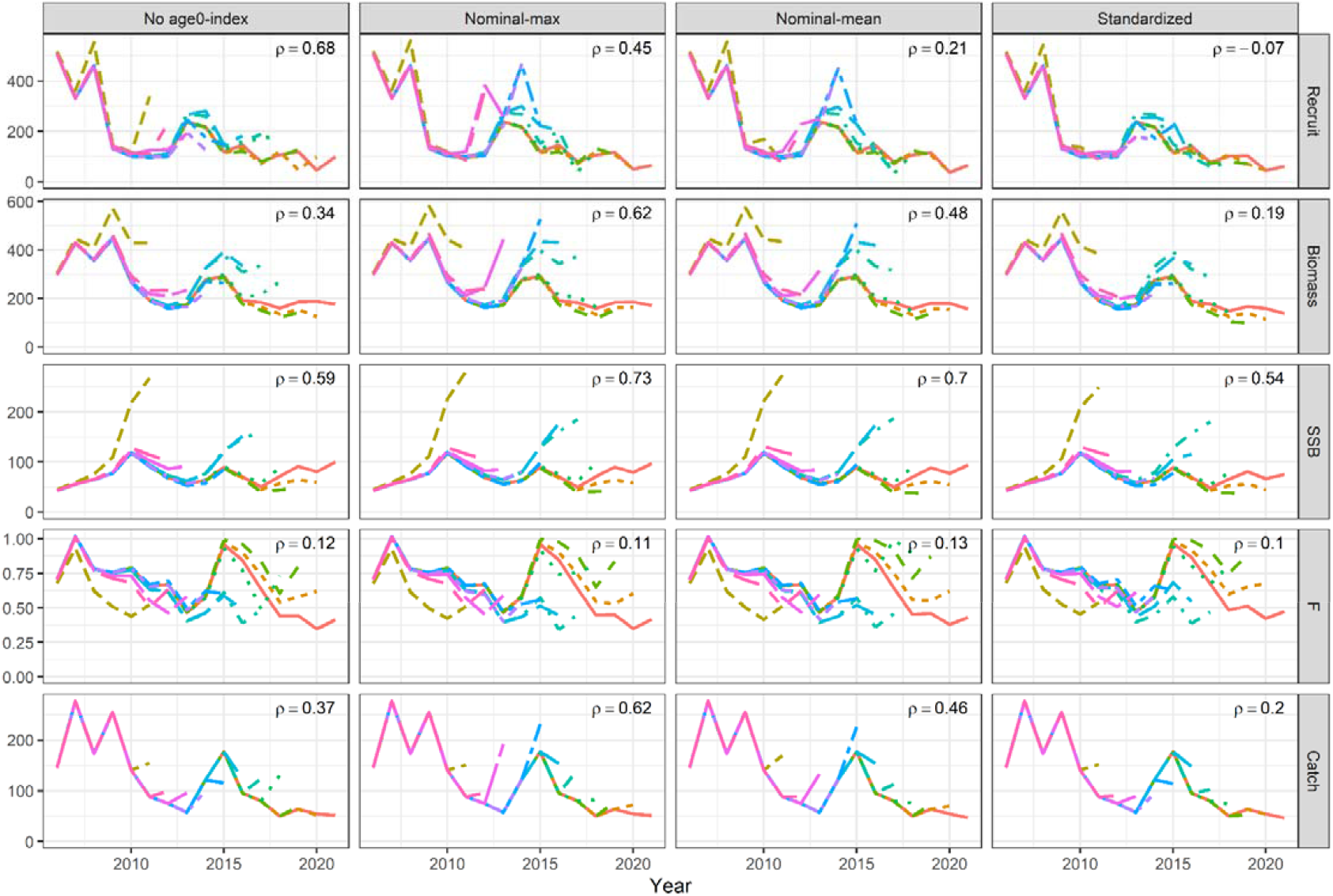
Results of retrospective forecasting for each VPA model. Mohn’s rho values are shown in the upper right corners.

## Discussion

We developed a method for the standardization of catch rates of juvenile pufferfish using surf-net survey data. Our approach described the pulse dynamics of juvenile fish by using a Gaussian function (or the quadratic function with a log link) and then investigated whether or not the parameters of the Gaussian function varied across years and should be estimated as fixed or random effects. We found that although AIC and BIC values favored different models in terms of types of time-varying parameters, LOOCV revealed that the model that estimated all time-varying parameters as random effects (selected by BIC) outperformed the model that estimated one of the time-varying parameters (peak size) as fixed effects (selected by AIC) (Table 1). We then evaluated the effectiveness of the standardized CPUE as the recruitment index of wild fish in the stock assessment model. We showed that the overestimation biases in retrospective forecasting were much lower in the model with the standardized abundance index than in the models with an unstandardized (nominal) index or without the age-0 abundance index (Fig. 6). Below, we discuss the effectiveness of our standardization approach and the importance of short-term forecasting in fisheries stock assessment and management.

### Effectiveness of our standardization approach

The most important point in our modeling is to use random effects in CPUE standardization, although the effects of years are usually estimated by fixed effects. This is because the surf-net survey data had a small sample size (5.83 samples per year, on average) and large observation errors. In particular, the total number of catches was zero in 2017; therefore, the estimate of the abundance index would be zero if using a fixed-effect model. This would be problematic in the stock assessment model because log-transformed values of abundance indices are usually used. This problem can be avoided by estimating time-varying parameters using random effects because the estimated variance in time-varying parameters (i.e., the magnitude of process errors) allows us to estimate expected CPUEs at appropriately positive values (Figs. 2, 4). Accordingly, our random-effect model could efficiently avoid overfitting, reduce uncertainty in observation errors, and smooth yearly trends in abundance indices. The availability of random effect models is growing in fisheries science along with the development and wide application of TMB (Nielsen and Berg 2014; Thorson and Barnett 2017; Nishijima et al. 2021). We demonstrate that a latent-variable model with random effects can provide an effective solution for CPUE standardization in data-poor situations.

To describe pulse patterns of expected catch rates of juvenile pufferfish, we used the Gaussian function or a quadratic function with a log link (Fig. 2). Although generalized additive models (GAMs) are often used to describe nonlinear responses of CPUE to a continuous variable, using GAMs in this study is not feasible owing to the small sample size. The Gaussian function is not flexible but easily describes the pulse dynamics of juvenile fish that utilize coastal sandy zones as short-term nursery grounds. The Gaussian function requires three parameters (Fig. 3), which can result in overparameterization when all three parameters are estimated as time-varying parameters. We therefore performed model selection based on information criteria and cross-validation in order to select a parsimonious model with high predictive ability. The model diagnostics showed no serious issues, such as residual trends, in our modeling (Online Supplementary Material). Our approach with the Gaussian function is a reasonable way to robustly estimate expected CPUEs and a standardized abundance index.

Using the Gaussian function provides insight into the biology of juvenile pufferfish because the estimated parameters are interpretable. The magnitude of peaks was estimated by a random walk, reflecting a gradual decrease rather than a random fluctuation in recruitment from wild fish (Fig. 3c). Although the cause of the decreasing trend is unknown and requires further investigations, this observation suggests that a sudden emergence of an unexpectedly strong year class is unlikely to occur. The timing of peaks was time-invariant (Fig. 3b), whereas the variation in the width of pulses varied independently (Fig. 3a). The data points for 2006 and 2009 revealed a delay in peak timing; however, peak timing showed high robustness overall (Fig. 2). The pulses in 2006 and 2015 were prolonged (Fig. 3a), which may reflect a phenological shift in arriving and leaving the nursery area. However, the changes in these two parameters were not unidirectional, suggesting that pufferfish are unlikely to show a phenological response in early life stages (between hatching and the juvenile stage) to climate change, including ocean warming, at present.

### Importance of short-term forecasting

We used retrospective forecasting to evaluate the effectiveness of CPUE standardization. The ‘usual’ retrospective analysis, which does not forecast future dynamics, is often conducted to compare stock assessment models with different abundance indices (Cao et al. 2017; Hashimoto et al. 2019). However, the aim of this study was to use survey data for juvenile pufferfish as prior information just before the annual stock assessment and fisheries recruitment; the survey is conducted from May to July, the stock assessment is conducted in August, and the age-0 fish begin to be caught by small-scale trawl fisheries in the autumn. We therefore performed retrospective forecasting to compare projections of population dynamics for the following year among models with the nominal-max index, nominal-mean index, standardized index, or no age-0 index. The overestimation bias was much higher for models with no age-0 index, the nominal-max index, and the nominal-mean index than for the model with the standardized abundance index (Fig. 6). The model without the age-0 index forecasted recruitment for the next year based on the latest 5-year average and resulted in the overestimation of recruitment because it did not account for the gradual decrease in recruits. The overestimation bias in recruitment was slightly reduced using nominal indices but was still large because the high variance in fitting of the abundance index (Fig. S3 in Online Supplementary Material) led to overestimated recruitment in a few years. The standardized index showed the smallest variance in fitting (Fig. S3 in Online Supplementary Material), stabilizing and improving the short-term forecasting of population dynamics. Although overestimation bias still remained for SSB, this bias was derived from a few years (2011, 2016, and 2017) and improved by using the standardized index. The bias in SSB might be explained by errors in age decomposition and, therefore, further improvements are not within the scope of this study. The importance of recruitment forecasting towards sustainable fisheries management is increasing under climatic and environmental variability (Beveren et al. 2021) and our study demonstrates the effectiveness of retrospective forecasting depending on model configurations.

The high predictive value of the model with the standardized index might be due to the use of VPA in the stock assessment model. We used VPA because it has been used for age-structured stock assessment for a long time in Japan (Ichinokawa et al. 2017). Although the state-space stock assessment model (SAM) is an alternative age-structured model (Nielsen and Berg 2014), it does not always outperform VPA with respect to estimation accuracy (Okamura et al. 2018). VPA employs the backward calculation of stock dynamics and, therefore, requires the post hoc calculation of future dynamics under an ad hoc assumption for recruitment (e.g., the latest 5-year average). By contrast, statistical catch-at-age models, such as SAM, based on forward calculation can estimate the recruitment process using the stock-recruitment relationship or random walk recruitment (Nielsen and Berg 2014; Okamura et al. 2018), which may diminish the need for an age-0 index for one-year forecasting. However, the stock of Japanese pufferfish around Ise-Mikawa Bay has an ambiguous stock-recruitment relationship with large uncertainty (Suzuki et al. 2021) and the simple random walk method (in which the number of recruits in a year is expected to be equal to that in the previous year) would likely overestimate the future number of recruits considering the decreasing trend. Because the standardized age-0 index based on the juvenile survey is a unique toolkit at present to determine the relative number of recruits since before pufferfish are caught by fisheries, the effectiveness of CPUE standardization for the age-0 index is not likely to change, even if using other stock assessment models.

One-year forecasting using the standardized recruitment index is expected to be advantageous for the stock assessment and management of the Japanese pufferfish over alternative approaches. Previous studies have shown that the effectiveness of short-term forecasting or reducing the data management time-lag can be large (Wiedenmann et al. 2016; Li et al. 2016) or not large (Zimmermann and Enberg 2017; Hutniczak et al. 2019; Okamura et al. 2020), depending on the characteristics of species, fisheries, forecasting accuracy, management objective, and survey costs. In the case of this pufferfish stock, young fish are caught mainly by longline fisheries and the number of recruits has gradually decreased, leading to the overestimation of ABC (Suzuki et al. 2021). Our one-year forecasting can reduce the risk of overfishing derived from the overestimation of ABC. Moreover, reducing the uncertainty and inaccuracy of stock assessment allows for the setting of a higher ABC (smaller ‘buffer’ in a harvest control rule) while reducing the probability of overfishing (Punt et al. 2012; Mildenberger et al. 2021). Our approach is expected to simultaneously reduce the risk of overfishing and increase sustainable yields. The use of environmental variables has been a major focus of research aimed at projecting future recruitment (Costello et al. 1998; MacKenzie et al. 2008); however, it is generally unlikely to be successful owing to the complex relationships between life history traits in early stages and environmental factors (Haltuch et al. 2019). Although incorporating recent information on fish abundance itself into stock assessments and future projections is promising, frequent and large-scale surveys are often labor-intensive and are not always cost effective (Zimmermann and Enberg 2017; Hutniczak et al. 2019). In addition, the availability of survey data has been limited to major fisheries stocks (Hashimoto et al. 2019). This paper provides a valuable case study demonstrating that accurate short-term forecasting of fish population dynamics is possible if small-scale survey data capture the relative abundance of fisheries stocks with simple life histories and limited distributions and are appropriately handled by latent-variable modeling. Even if the sample size is small, the use of dormant, unused data in an appropriate statistical framework, such as latent-variable modeling, will contribute to reliable stock assessment and sustainable management in many fisheries resources.

## Acknowledgements

We thank the members of Mie Prefecture Fisheries Research Institute who participated in the surf-net survey. This study was funded by JSPS KAKENHI Grant Number 19K15905. This study was partially supported by grants for marine fisheries stock assessment and evaluation in Japanese waters from the Fisheries Agency and the Fisheries Research and Education Agency of Japan.

