## Supplementary material for "Modeling Pulse Dynamics of Juvenile Fish Enables the Short-term Forecasting of Population Dynamics in Japanese Pufferfish: A Latent Variable Approach": Table A1

Table A1: Information on surf-nets

| Year | Wind-net length (m) | Wind-net height (m) | Mesh size (mm) | Bag-net length (m) |
| --- | --- | --- | --- | --- |
| 2004-05 | 5.0 | 1.5 | 0.5 | 3.0 |
| 2006-07 | 10.0 | 1.0 | 2.7 | 2.0 |
| 2008-21 | 4.0 | 1.0 | 1.0 | 3.0 |
