## Supplementary material for "Modeling Pulse Dynamics of Juvenile Fish Enables the Short-term Forecasting of Population Dynamics in Japanese Pufferfish: A Latent Variable Approach": Online Supplementary Material

**Model Diagnostics for CPUE Standardization**

We conducted model diagnostics to investigate whether CPUE standardization was appropriately performed. We used a negative binomial distribution for the discrete variable (i.e., integer larger than or equal to zero). In such a case, a simulation-based approach can be effective (Hartig 2020). We used the R package ‘DHARMa,’ which simulates readily interpretable scaled (quantile) residuals for fitted generalized linear mixed models (Hartig 2020). This approach first simulates bootstrapped data for response variables under the assumption that the fitted model is true. Standardized residuals are obtained using the function ‘createDHARMa(),’ which calculates quantiles of observed samples against cumulative probability distributions computed from simulated samples. The standardized residuals have a value from zero to one and ideally follow a uniform distribution within this range. Because simulated samples have the same values when a discrete variable is used, as in this study, random variables are added from the uniform distribution from −0.5 to 0.5 to both observed and simulated samples. In this way, we simulated a set of catch numbers 10,000 times from predicted CPUEs (equations 1 and 2 in the main text) and dispersion parameter 𝜑 in the best model and then calculated standardized residuals for each sample.

We performed the Kolmogorov–Smirnov test to examine whether the standardized residuals deviate from the uniform distribution and generated a QQ plot (indicating the relationship between theoretical quantiles and standardized residuals). We also conducted dispersion test, the outlier test, and the zero-inflation test, all of which are implemented in the package ‘DHARMa’ (Hartig 2020). We then analyzed the relationship between standardized residuals and predicted catch numbers. The predicted numbers were rank-transformed and then scaled from zero to one as in Hartig (2020). We fitted a generalized additive model (GAM) using the function ‘gam()’ in the R package ‘mgcv’ (Wood 2011). We used a beta distribution with a logit link because the response variables (i.e., the standardized residuals) ranged from zero to one. In addition, we analyzed the relationship between standardized residuals and the explanatory variables (years and time since May 1). Although we show the case in which non-transformed explanatory variables are used, similar results are obtained when using rank-transformed explanatory variables. We also applied GAM by treating the explanatory variables of year and time since May 1 as continuous.

The Kolmogorov–Smirnov test showed no significant deviation in standardized residuals from the uniform distribution (*p* = 0.248, Fig. S1a). The Q-Q plot also showed little deviance from the theoretical prediction (Fig. S1b). Both the dispersion and outlier tests showed no significant effects (*p* = 0.934 in the dispersion test and *p* = 1.00 in the outlier test). Although the number of observed zero samples (39) was slightly lower than the median number of simulated zero samples (43), it did not deviate significantly from the theoretical expectation (*p* = 0.362, Fig. S1c). This suggests that it is not necessary to apply a zero-inflated model to the dataset. There were no significant relationships between standardized residuals and catch numbers (*p* = 0.359, Fig. S1d), year (*p* = 0.720, Fig. S1e), or date (*p* = 0.814, Fig. S1f). In total, the function and probability distribution in the best model we selected were well suited to the observations.

Moreover, we performed a retrospective analysis to check the robustness of the standardized index by sequentially removing the latest years (10 years at maximum). The past estimates did not change substantially by removing recent data (Fig. S2), supporting the robustness of the standardized index against data updating.

We also conducted model diagnostics for the fitting of the tuned VPA to the nominal max, nominal mean, and standardized indices. The standard deviation of fitting was much smaller for the standardized index than for the nominal indices (Fig. S3). GAM showed that the trends in residuals were not significant for all three indices. Considering all model diagnostics together, we did not detect any serious issues with the standardized index.

**Figures**

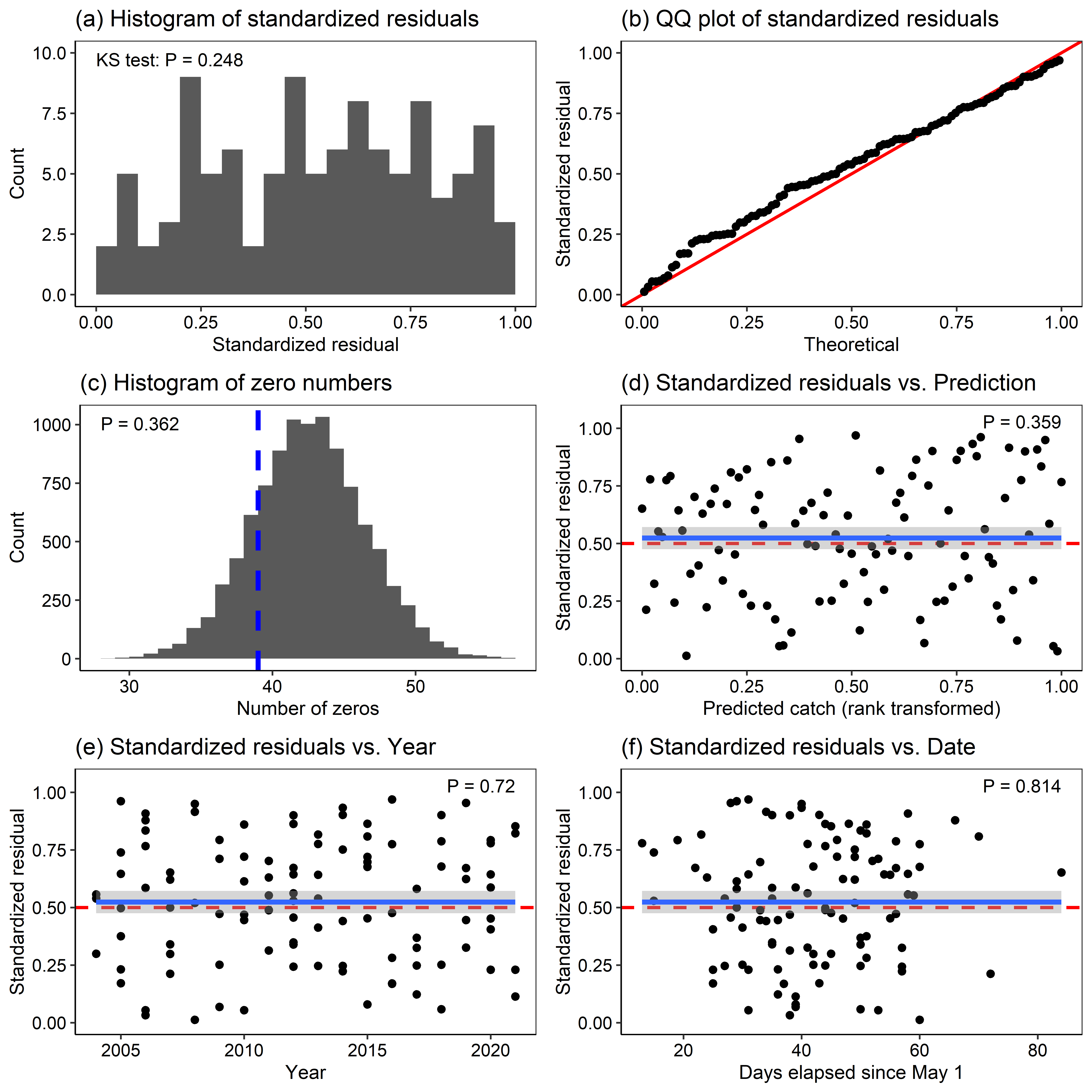
Figure S1: Results of model diagnostics. (a) Histogram of standardized residuals, (b) QQ plot of standardized residuals and theoretical quantiles, (c) a histogram of the numbers of simulated zero samples, (d) the relationship between standardized residuals and (rank-transformed) predicted catch numbers, (e) the relationship between standardized residuals and years, and (f) the relationship between standardized residuals and days elapsed since May 1. Red lines in b and d–e represent theoretical expectations. The blue dashed line indicates the observed number of zero samples. Blue lines in d–f show estimated regression lines by GAMs with 95% confidence intervals.

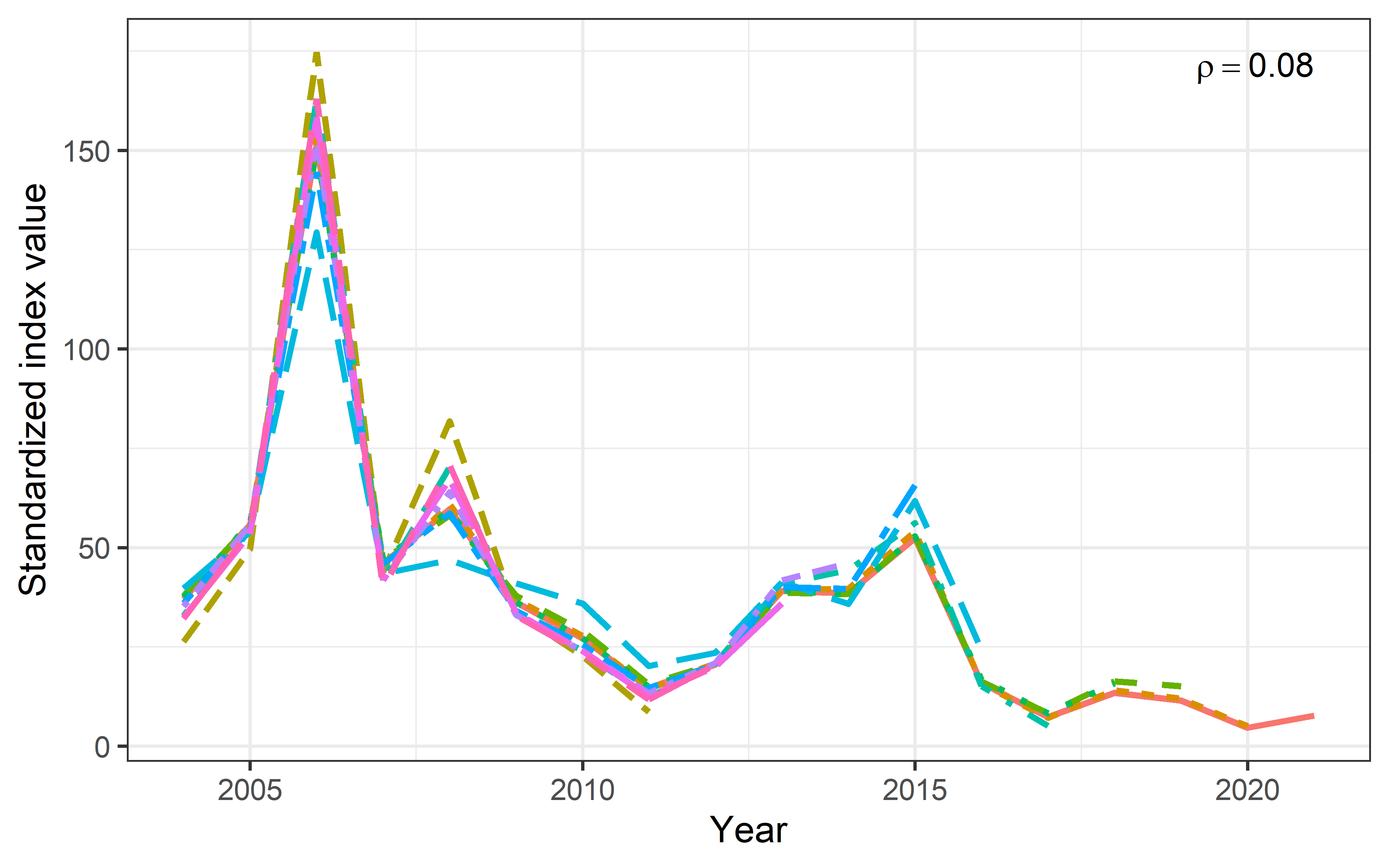

Figure S2: Retrospective patten for the standardized index. Mohn’s rho value is shown in the upper right corner.

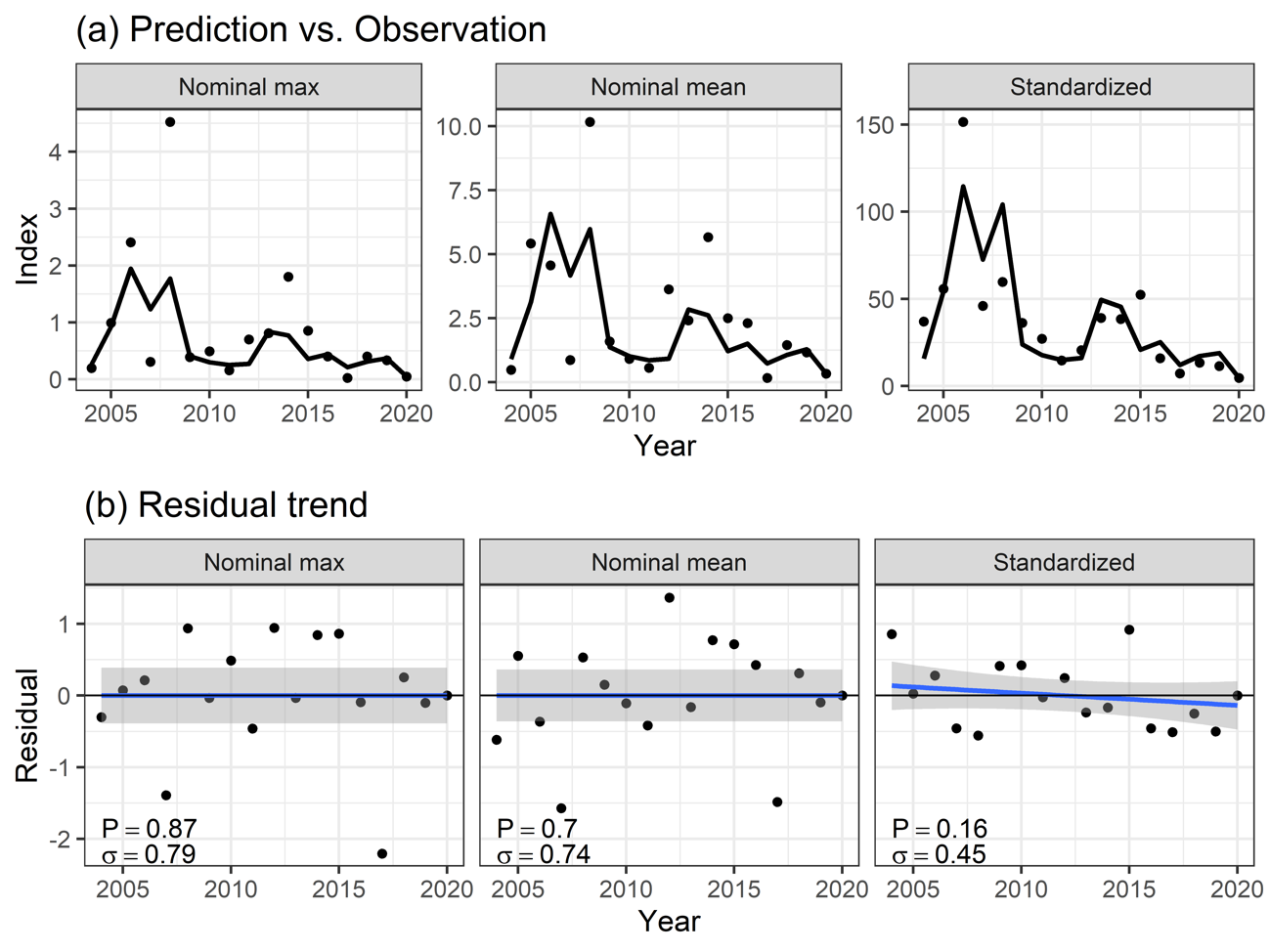

Figure S3: Fitting of abundance estimates to the nominal max index (left), the nominal mean index (middle), and the standardized index (right). (a) Predicted values (lines) and index values (dots). (b) Trend in residuals. Blue lines indicate predicted regression lines by GAM with 95% confidence intervals. *p*-values and standard deviations are shown in the lower-left corners.
